# Kinetic and thermodynamic analysis of PKA-R–cAMP interactions in crude media using focal molography

**DOI:** 10.1101/2025.10.01.679885

**Authors:** John Oehninger, Daniela Bertinetti, Friedrich W. Herberg, Andreas Frutiger

## Abstract

Quantifying biomolecular interactions under near-physiological conditions is essential for understanding biological processes. However, this remains a challenge in established techniques: for surface plasmon resonance, bulk refractive-index shifts and non-specific adsorption in hydrogel matrices can dominate the signal in serum, whereas for isothermal titration calorimetry, complex media primarily introduce large, composition-dependent background heats and run-to-run variability that obscure the binding enthalpy. Here, we demonstrate that focal molography, a label-free optical biosensing method, can reliably measure the kinetics and thermodynamics of the protein kinase A regulatory subunit binding to cyclic AMP derivatives in both buffer and 50% human serum. By performing the kinetic measurement over a range of temperatures, we were able to reveal that binding remains highly favorable in both environments, yet the results suggest that the balance of driving forces shifts toward enthalpic contributions in serum. This hints at the substantial difference of interaction mechanisms in a complex biological media compared to buffer systems. Our findings show that focal molography reliably quantifies protein-ligand interactions in complex media, providing consistent kinetic and thermodynamic data while overcoming limitations of existing methods. This highlights the potential of focal molography as a valuable tool for studying interactions under near-native conditions.

## Introduction

Understanding biomolecular interactions and drug behavior in physiologically relevant environments is critical for decoding cellular processes and developing effective therapeutic interventions. Traditionally, interactions are studied and drugs are screened in purified buffer systems, which fail to replicate the complexity of physiological environments [1]. These systems overlook essential factors such as competing biomolecules, enzymatic degradation, reduced effective concentrations, and macromolecular crowding, all of which can significantly influence binding kinetics and drug stability *in vivo* [2–5]. Studying these interactions early in the drug development process can improve predictive accuracy, reduce costly late-stage failures, and accelerate the path to viable therapeutics [6]. Even beyond these broader physiological differences, buffer systems themselves can introduce misleading artefacts. Buffers often contain stabilizing agents, salts, or chelators that interact with proteins or ligands in unintended ways—potentially masking binding sites, altering conformational equilibria, or stripping essential co-factors. Indeed, even common buffers such as HEPES, MES, Tris, and phosphate have been shown to significantly affect the thermal stability and substrate-binding activity of RecA in a concentration-and pH-dependent manner [7]. Likewise, reviews confirm that buffer composition and ionic strength can alter both equilibrium and kinetic binding constants in drug assays [8–10]. This tightly controlled pH and absence of reactive species or enzymatic activity in such systems may stabilize compounds that would otherwise degrade rapidly in biological fluids, leading to an overestimation of drug stability and potency. As a result, buffer-based assays can produce overly clean kinetic profiles that fail to capture multiphasic or cooperative binding behavior observed in complex media [5].

Despite their widespread adoption in biophysics labs, conventional techniques face different serum-related limitations. Surface Plasmon Resonance (SPR) is highly sensitive to bulk refractive index changes, temperature drift, and non-specific adsorption within carboxymethyl-dextran matrices, which makes serum injections particularly prone to artefacts and referencing mismatches [11]. Isothermal Titration Calorimetry (ITC), by contrast, is performed in solution and is less affected by surface fouling, its main limitation in serum is the large, composition-dependent heat background (and biological variability across lots) that is difficult to subtract accurately with controls, making binding enthalpies hard to interpret.

Recently a label-free optical method termed focal molography was proposed to enable more physiologically relevant biomolecular interaction analysis. Focal molography employs a two-dimensional sub-micron pattern of molecular recognition sites in the shape of a diffractive lens, termed a *mologram* [12, 13]. The pattern is composed of almost a thousand 200 nm lines (the ridges) and referencing areas (the grooves) in between. Light is diffracted by the molecules that bind to the pattern and interferes constructively in the case of the ridges and destructively in the case of the grooves in a focal point. The light intensity diffracted is quadratically proportional to the difference in the number of molecules in ridges and grooves. This diffraction-based detection enables precise, real-time monitoring of molecular interactions with high sensitivity and robustness in physiologically relevant systems [12, 14–16].

The key distinction between focal molography and refractometric techniques lies in its distributed, sub-micron-scale self-referencing mechanism and its detection of only the differential signal at the focal point [17, 18]. In contrast, refractometric sensors like SPR rely on a macroscopically separated reference channel—often millimeters away—and measure signal and reference independently. As the distance between signal and reference regions increases, it becomes increasingly difficult to ensure that non-specific effects, such as bulk refractive index changes or temperature shifts, affect both regions identically (are correlated on both regions). By performing referencing at a molecular length scale, focal molography overcomes this challenge, delivering robust, drift-free performance even in complex media such as human serum, where traditional sensors often fail due to interference from non-specific effects [19]. By canceling bulk effects at the micrometer scale, focal molography avoids the refractometric serum artefacts typical of SPR, and because it reads differential coherent mass rather than total heat, it is not confounded by the high baseline heats that complicate ITC.

In this study, we use focal molography to investigate the interaction between the cAMP binding domain A of the bovine R subunit I*α* of PKA (bRI*α* 92–260) (PKA-R) and cyclic AMP (cAMP) analogues in both buffer and 50% human serum. The R-subunit inhibits protein kinase activity of the catalytic subunit of PKA by direct binding. By quantifying kinetic and thermodynamic parameters across a physiological temperature range, we examine how the biological environment alters the enthalpic and entropic contributions to binding. We compare our findings to established SPR and ITC data, demonstrating that focal molography enables robust quantification of protein-ligand interactions under near-physiological conditions.

## Materials and methods

### Materials

To prepare the sensors for interaction studies, we immobilized cAMP derivatives covalently on the chip surface. We selected derivatives that feature a primary amine, which allowed for the stable conjugation of the ligands to the chip’s surface and derivatives that were shown previously to maintain their biological activity when linked to a surface [20]. The 6-AH-cAMP [N^6^-(6-Aminohexyl)adenosine-3*^′^*,5*^′^*-cyclic monophosphate] and 8-AHA-cAMP [8-(6-Aminohexylamino)adenosine-3*^′^*,5*^′^*-cyclic monophosphate] derivatives were obtained from Biolog Life Science Institute (Bremen, Germany). Their molecular structure is shown in Fig 1. Additional reagents for buffer preparation and surface immobilization were obtained from Sigma-Aldrich.

**Fig 1.**
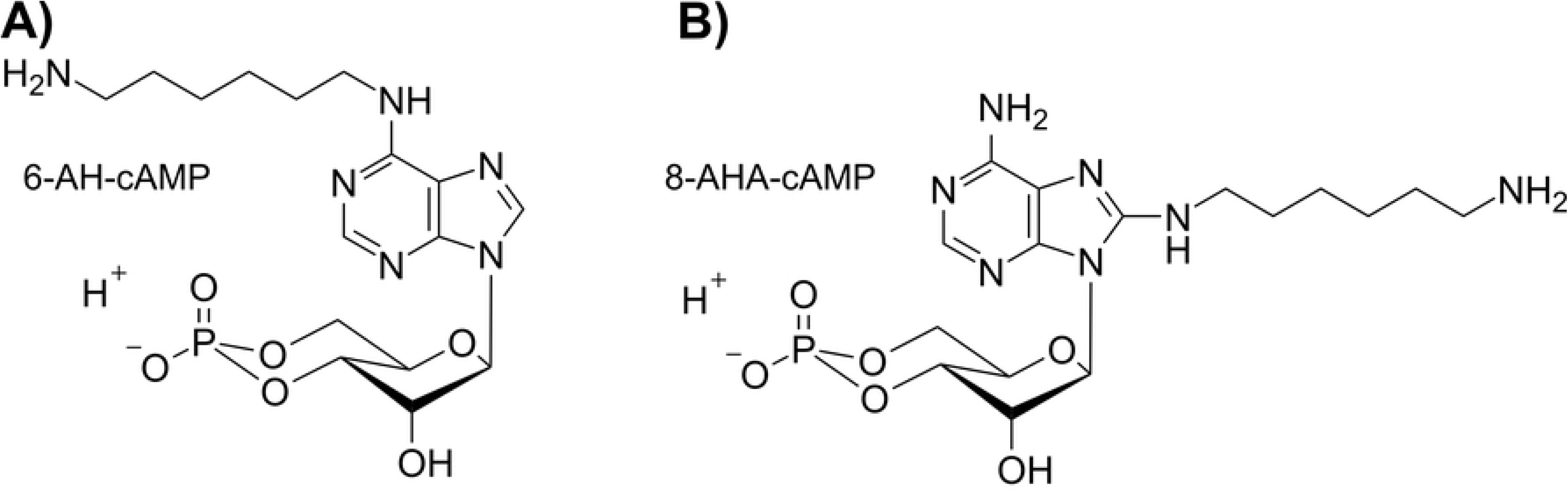
Molecular structures of the two cAMP derivatives used in this study. **(A)** 6-AH-cAMP features an aminohexyl group at the N6 position of the adenine base, while **(B)** 8-AHA-cAMP contains an aminohexylamino modification at the 8th position for covalent immobilization. Both derivatives retain the characteristic cyclic phosphodiester bond of cAMP, ensuring biological activity. These derivatives were used as ligands for PKA-R binding studies in this work.

The recombinant bovine RI*α* 92-260 protein (19,035.5 Da), a truncated form of the regulatory subunit of cAMP-dependent protein kinase A (PKA-R), was used in all experiments. This construct retains the essential cAMP-binding domain (residues 92–260), facilitating 1:1 stoichiometry with cAMP derivatives while simplifying data analysis compared to the full-length multi-domain protein, and is hereafter referred to as PKA-R [21]. The cDNA encoding RI*α* 92-260 was expressed in *E. coli* BL21 (DE3) RIL cells, and the protein was purified to obtain a nucleotide-free state following the procedure described in Moll et al. (2007) [20] and Hahnefeld et al. (2005) [22]. Briefly, the protein was eluted from 6-AH-cAMP agarose using 8 M urea in 20 mM MOPS (pH 7.0), followed by extensive dialysis against 150 mM NaCl, 20 mM MOPS, 10 mM MgCl_2_, and 1 mM ATP (pH 7.0). Protein purity was confirmed by SDS-PAGE, and its biological activity was verified using a phosphotransferase assay with Kemptide as substrate. Protein concentration was determined using a Bradford assay. All chemicals and reagents used were of analytical grade or higher.

### Instrumentation

Focal molography experiments were conducted using a prototype instrument from the“Callisto Pre-series Generation” developed by lino Biotech AG (Adliswil, Zurich, Switzerland). The custom-built device is equipped with three data generation channels that can be used to characterize the molecular interaction of interest, a molographic detection channel (excited by a 785 nm laser) which measures the focal spot intensity of the mologram, a refractometric detection channel, which measures the shift of the spot in the focal plane, as well as a fluorescence microscopy detection channel (660 nm excitation wavelength) in TIRF mode [12, 23]. Both lasers are coupled into a dielectric slab waveguide made from Ta_2_O_5_ as described earlier [12, 13]. The instrument is compatible with multiple flow chamber configurations, allowing for single as well as quadruple flow chambers. The chambers are made from PEEK (polyetheretherketone) polymer, with a support made from black anodized aluminum and flat seals laser-cut from Kalrez sheets (DuPont). The quadruple flow chambers each cover one row of molograms (9 per row). All experiments were carried out using the quadruple flow chamber, in which each flow channel has a width of 1 mm, a height of 100 µm and a length of 12 mm, providing optimal flow conditions for interaction studies. The flow chamber assembly process is shown in Fig 2C and D. The sensor chip, mounted within the chamber (see Fig 2D), measures 9 mm x 18 mm and features square sensor spots measuring 400 µm x 400 µm arranged in a 6 x 9 grid (54 molographic measurement spots in total). The distance between sensor spots in a row is 690 µm and the spacing between rows is 750 µm. The sensor chip features an in-and out-coupling grating that are used to couple light in and out of the waveguide as described earlier [13]. Coherent mass density quantification is performed from the outcoupled light power according to the models described in [24], refractometric mass density quantification is performed according to the models described in [23]. For automated sample and buffer delivery, an Alias Autosampler (Spark Holland, Emmen, Netherlands) was employed in combination with an external pump (Hamilton PSD/4 Stand-alone Pump) with a 2.5 mL syringe (UHMWPE plunger material). The autosampler had an internal 6-channel selection valve with 6 connections going from the sampler to the chip of which 4 were used for the 4-channel flow chamber. Temperature control was maintained using an external liquid heater/chiller (Thorlabs, LK220). The liquid coolant lines are connected to a circulation system that runs through the chip mount stage, which is made of black anodized aluminum, allowing efficient heat transfer and distribution throughout the mounting plate. This ensured precise temperature adjustments throughout the experiments with a precision of ± 0.1 K. Data acquisition and analysis were performed using proprietary software developed by lino Biotech AG. A detailed image of the complete experimental setup can be seen in Fig 2A.

**Fig 2.**
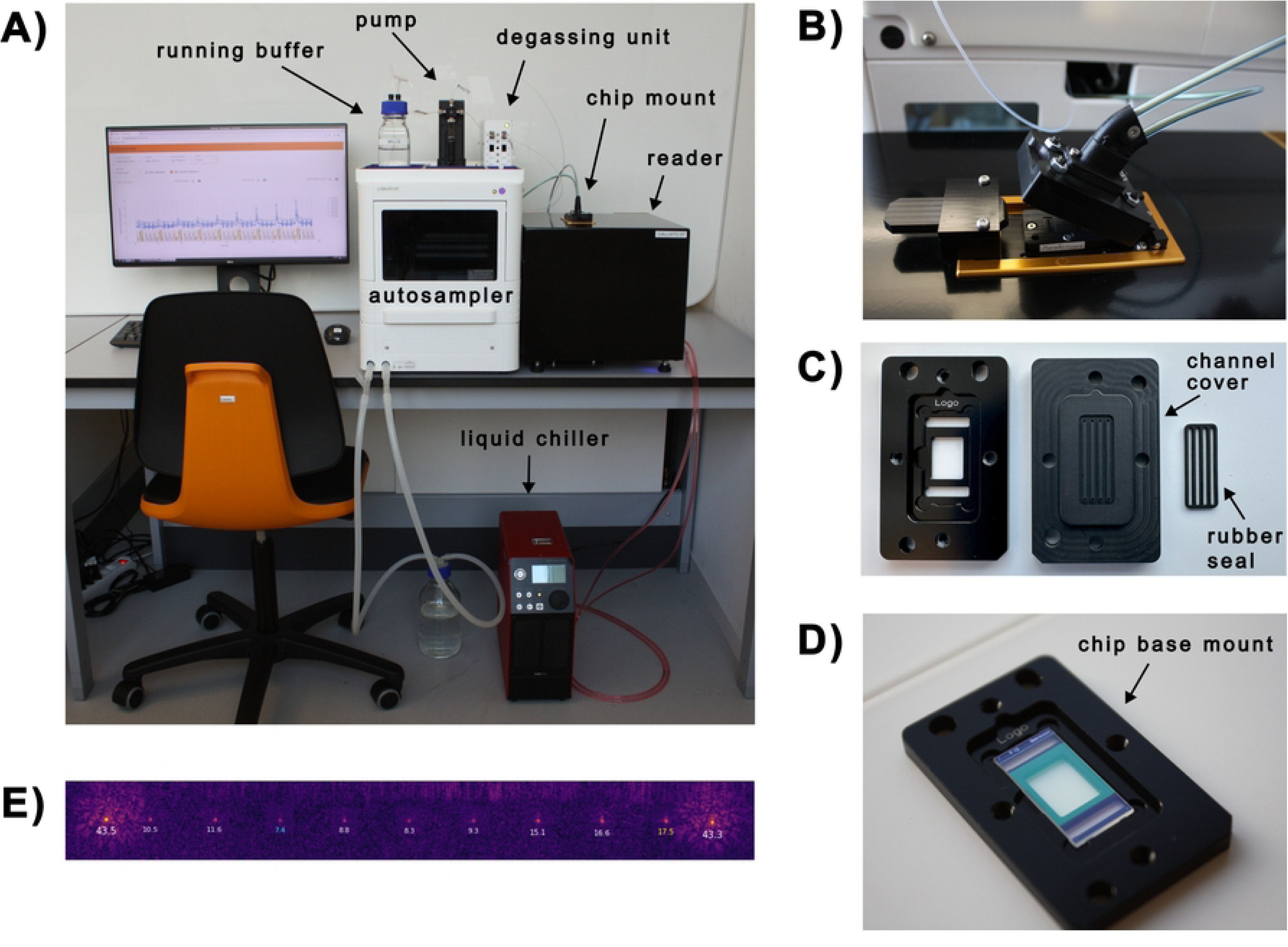
Instrumentation setup for focal molography experiments. **(A)** Overview of the complete experimental setup, including the reader instrument, autosampler, pump, degassing unit, chip mount, and liquid chiller for precise temperature control. **(B)** Close-up of the chip mount with connected fluidic lines. **(C)** Disassembled view of the flow chamber assembly, showing the aluminum chip base mount (left), PEEK flow chamber with integrated channels (middle), and laser-cut Kalrez rubber seal (right). **(D)** Aluminum chip base mount with the sensor chip mounted. **(E)** Representative molographic readout image from the instrument showing diffracted signal spots along a measurement row (9 spots). The two molograms on the left and right are reference molograms.

### Sensor chip preparation, conjugation and immobilization

Sensor chip preparation involved three key steps: fabrication of molograms via reactive immersion lithography (RIL), chemical conjugation of cAMP derivatives to linker molecules, and subsequent immobilization of the conjugates onto the sensor surface.

### Mologram fabrication via reactive immersion lithography (RIL)

Each sensor chip features 54 molograms fabricated via RIL [12, 25]. This process uses a dielectric waveguide surface coated with photoactivatable polyacrylamide-grafted polyethylene glycol copolymer brushes (PAA-g-PEG-NH-PySNPPOC, SuSoS AG, Switzerland). The brush layer forms a non-fouling background that suppresses non-specific binding while preserving functionality of the immobilized biomolecules. [12].

A slight variation to the published protocol for the photolithography was employed: a more hydrophilic pyridine group was used as the photoprotective moiety of the primary amines (PySNPPOC - Carbamic acid 2-[4-ethyl-2-nitro-5-(pyridin-2-ylsulfanyl)phenyl]propyl ester] purchased from Orgentis Chemicals, Germany). Molograms were patterned by photolithography using a phase mask, which selectively removed the protective groups in the ridges, creating a line pattern of exposed primary amino groups. These amines were functionalized with 1 mM MeTz-PEG4-NHS (Methyltetrazine-(polyethylene)-N-hydroxysuccinimide, Vector Laboratories, USA) in HBS-T buffer (50 mM HEPES, 150 mM NaCl, pH 8.0) for 1 hour, then washed sequentially with DMSO, isopropanol (IPA), and Milli-Q water.

Following a flood exposure step to deprotect the remaining regions [13], the grooves were passivated with 10 mM MeO-PEG2-NHS (Broadpharm, USA) in HBS-T (pH 8.0). This resulted in spatially separated regions containing clickable MeTz-groups and PEG-passivated amines, embedded in a two-dimensional layer of the underlying PAA-g-PEG polymer roughly 10–20 nm in thickness in the hydrated state. Mologram architectures are denoted as [ridges|grooves]; the resulting structures are therefore referred to as [MeTz|PEG] molograms.

### Conjugation of cAMP derivatives

To enable site-specific immobilization, the cAMP derivatives 6-AH-cAMP and 8-AHA-cAMP were conjugated to TCO-PEG4-TFP linkers (Transcyclooctene-polyethyleneglycol-tetrafluorophenyl, Vector Laboratories, USA) via amine-reactive coupling. A 5:1 molar excess of cAMP derivative to TCO linker was used to drive the reaction to completion and avoid the need for purification.

Specifically, 200 µL of a 5000 µM cAMP derivative solution in DMSO (dimethylsulfoxide) was mixed with 2 µL of 100 mM TCO-PEG4-TFP (in DMSO) and 198 µL of HBS-T buffer (pH 8.0). The reactants were incubated for 1 hour at 15*^◦^*C on a shaker at 600 rpm to ensure efficient conjugation between the TFP esters and primary amines on the cAMP derivatives.

### Ligand immobilization onto sensor surface

The resulting cAMP-TCO conjugates were applied directly to the sensor surface for immobilization via MeTz-TCO click chemistry [26]. A 100 µL aliquot of the conjugate solution was pipetted onto the chip and incubated for 1 hour at room temperature.

After incubation, the chip was sequentially washed in NaOH, toluene, isopropanol, Milli-Q water, and 3 M guanidine hydrochloride (GuHCl), with each wash lasting 30 seconds on a shaker. The chip was then dried with nitrogen gas and stored at 4*^◦^*C in an Eppendorf tube until use.

Immediately before each experiment, the chip was rinsed with Milli-Q water, dried with inert gas, and mounted into the flow chamber for measurements.

### Experimental setup and procedure

All interaction analyses were conducted in a buffer, composed of 150 mM NaCl, 20 mM MOPS, 10 mM MgCl_2_, 15 µM BSA and 0.005% (v/v) Tween-20 at pH 7.0 (termed *buffer A*). Importantly, the BSA merely served as a blocking agent to reduce the sticking of the PKA-R protein to the sensor surface and the buffer was not matched in refractive index to the used human serum. For experiments conducted in human serum, a 50% dilution of thyroid-stimulating hormone (TSH) depleted and processed human serum (from Veritas Innovation LLC, Austin, TX, USA) in *buffer A* was used. Both buffer and serum experiments followed the same protocol. It is important to note that during the dissociation phase, *buffer A* was consistently used for both buffer and serum experiments, which was supplemented with BSA. The standardized sequence of injections, including durations and flow rates, is summarized in Table 1. Additionally, regeneration was performed three times to ensure complete removal of bound analyte.

**Table 1.** Injection sequences used in all experiments, detailing each step’s sample type, duration, and flow rate.

| Type | Sample | Duration | Flow rate |
| --- | --- | --- | --- |
| Baseline | Buffer A | 1 min | 30 $\mu\text{L}/\text{min}$ |
| Association | PKA-R | 1 min | 30 $\mu\text{L}/\text{min}$ |
| Dissociation | Buffer A | 3 min | 200 $\mu\text{L}/\text{min}$ |
| Regeneration | 3 M GuHCl | 40 s | 300 $\mu\text{L}/\text{min}$ |
| Rinse | Buffer A | 1 min | 100 $\mu\text{L}/\text{min}$ |

The sensor chip was mounted in the aforementioned four-flow chamber system, which provided optimal fluidic conditions for kinetic measurements. As the flow chambers are addressed individually, only data from a single row of 9 molograms can be recorded at a time. Temperature control was maintained at 15*^◦^*C, 20*^◦^*C, 25*^◦^*C, 30*^◦^*C and 35*^◦^*C using the Thorlabs LK220 Thermoelectric Liquid Chiller.

### Data acquisition and kinetic analysis

Data was collected using the described instrumentation at maximum available sampling frequency which averaged to approximately 0.3 Hz. The instrument generated sensorgrams that were subsequently analyzed to extract kinetic and thermodynamic information using a dedicated kinetic fitting library provided by lino Biotech AG. This code leverages the widely-used Python module lmfit to perform nonlinear least-squares fits [27]. To analyze the binding data, a Langmuir 1:1 binding model was employed.

This model assumes a simple biomolecular interaction and fits three parameters *k*_on_, *k*_off_ and *R*_max_. The kinetic parameters extracted include the association rate constant (*k*_on_), dissociation rate constant (*k*_off_), and equilibrium dissociation constant (*K*_D_). The equilibrium dissociation constant was calculated using the following equation:

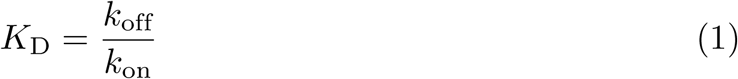

The fitting process was conducted globally, fitting all data points across different analyte concentrations simultaneously to enhance robustness and reliability.

Thermodynamic parameters, such as Gibbs free energy (Δ*G*), enthalpy (Δ*H*), and entropy (Δ*S*), were derived from van’t Hoff plots using temperature-dependent binding data. These parameters were calculated using the following relationships:

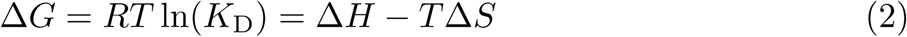

and

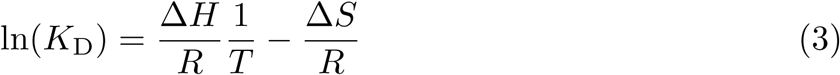

Here, *R* is the universal gas constant, and *T* is the absolute temperature in Kelvin. The slope of the van’t Hoff plot (ln(*K*_D_) vs. 1*/*T) provides the enthalpy change (Δ*H*), while the intercept gives the entropy change (Δ*S*).

### Calculation of transition-state thermodynamic parameters

The thermodynamic parameters of the transition state for the association 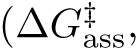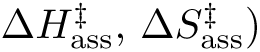 and dissociation 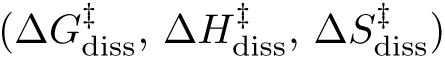 were calculated using kinetic rate constants (*k*_on_ and *k*_off_). According to transition state theory, the association rate constant *k*_on_ is given by:

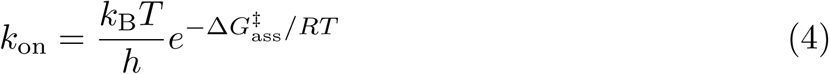

Here, *h* is the Planck constant and *k*_B_ the Boltzmann constant. After substituting in eq. 2 and taking the natural logarithm this results in a recasted Eyring equation in linear form:

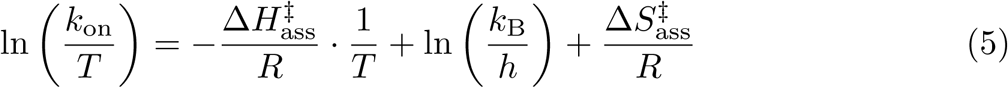

In this form, plotting ln (*k*_on_*/T*) against 1*/T* yields a line, where the slope is 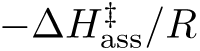 and the intercept is 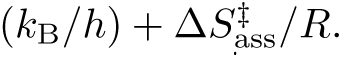 This allows the determination of both the enthalpy 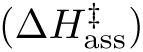 and entropy 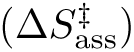 of activation. The same procedure applies to the dissociation rate constant *k*_off_.

### Controls and validation

#### Positive control: verification of specific PKA-R binding

To validate the successful immobilization of cAMP derivatives on the sensor chip, a positive control experiment was performed by injecting 320 nM PKA-R in *buffer A* over the sensor surface immobilized with the 6-AH-cAMP analog. This concentration was chosen based on prior studies to generate a strong response [20]. As shown in S1 Fig, the injection of PKA-R resulted in a sharp increase in coherent mass density (≈ 20 pg/mm^2^ within 60 seconds), indicating a strong and specific interaction with the immobilized cAMP. The signal remained stable over time, confirming that the observed response is due to specific PKA-R binding rather than non-specific adsorption.

#### Negative control: Competition with free cAMP results in no binding of PKA-R to the surface

To further confirm the specificity of the measured interaction, a negative control experiment was conducted using a pre-formed PKA-R–cAMP complex. A solution containing 160 nM PKA-R and 1.6 µM 6-AH-cAMP (1:10 molar ratio) was injected over the sensor chip to ensure that nearly all PKA-R binding sites were occupied by the ligand before exposure to the immobilized cAMP. As illustrated in S2 Fig, the injection of the complex did not produce an increase in coherent mass density, with only minor fluctuations (≈ 0.5 pg/mm^2^), consistent with baseline noise. This confirms that the focal molography sensor detects only free PKA-R molecules and does not register signals from PKA-R already complexed with cAMP. These results demonstrate that the sensor specifically detects PKA-R–cAMP binding events rather than non-specific interactions of the protein with the surface.

#### Rejection of non-specific binding in 50% TSH depleted human serum

A critical challenge when measuring biomolecular interactions in complex biological media is the potential for non-specific binding of serum components. To assess whether human serum produces a sensor response due to non-specific binding, a blank of 50% human serum (HS) in running *buffer A* was injected. It is important to emphasize that the running buffer is not refractive index matched to the serum, which would be required if a similar experiment were to be attempted with a refractometric sensor. As depicted in Fig 3, the injection of 50% HS alone did not result in a significant change in coherent mass density, indicating that the sensor does not detect non-specific adsorption from serum proteins. This confirms that binding signals observed in subsequent experiments arise from specific PKA-R–cAMP interactions rather than serum-induced artefacts.

**Fig 3.**
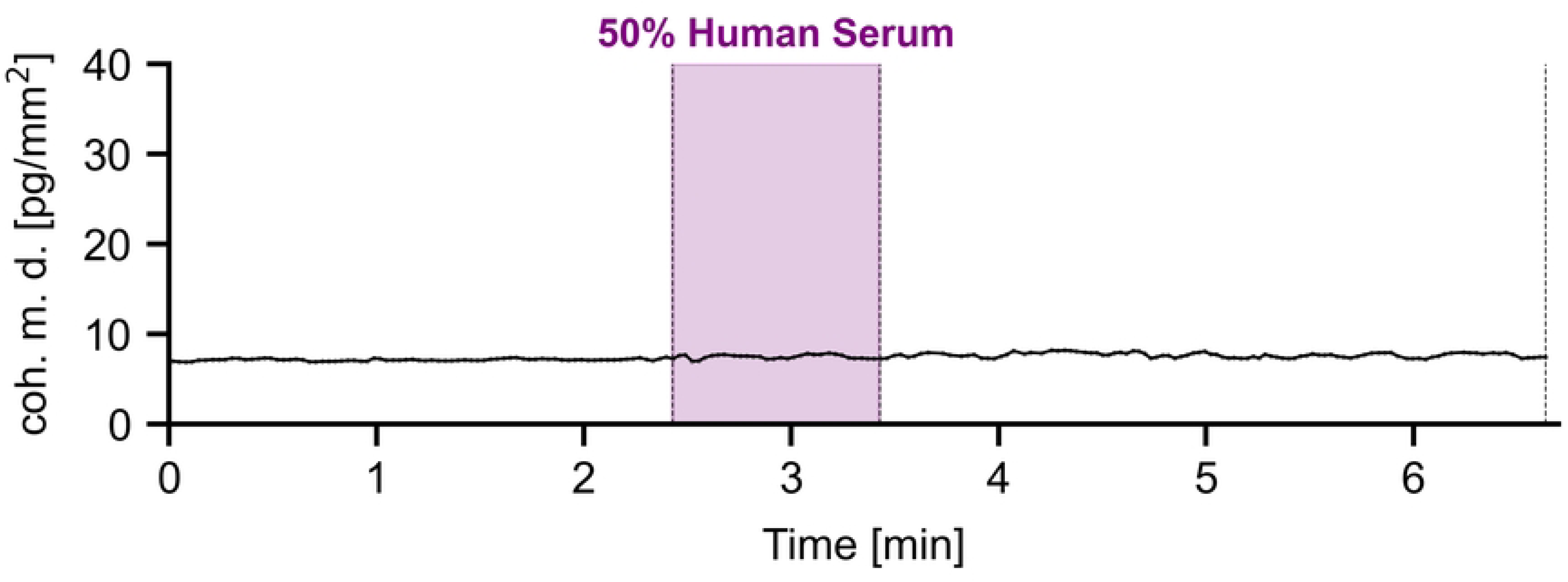
Control experiment demonstrating suppression of artefacts due to non-specific serum interactions. Injection of 50% human serum (HS) resulted in minimal signal change, indicating that the focal molography sensor effectively suppresses background noise from serum proteins.

## Results

### Overview of experimental analysis

This section is divided into two parts. The first part focuses on characterizing the interaction between PKA-R and cAMP derivatives in both buffer and 50% human serum, demonstrating the robustness of focal molography to assess kinetic parameters in complex biological environments.

The second part shifts the focus to the thermodynamic analysis of the interaction, exploring the temperature dependence of the binding events and calculating key thermodynamic parameters, including the Gibbs free energy (Δ*G*), enthalpy (Δ*H*), and entropy (Δ*S*). This analysis highlights subtle differences between buffer and serum environments, providing further insight into the behavior of PKA-R–cAMP interactions under different physiological conditions.

### Part 1: Characterization of PKA-R–cAMP interactions in buffer and serum

#### Kinetic measurements and titration series

In order to determine the kinetic parameters of the PKA-R–cAMP interaction, multi cycle kinetic experiments were conducted in both buffer and 50% human serum at 20*^◦^*C with intermediate regeneration. PKA-R concentrations ranged from 2.5 nM to 640 nM and intermediate regeneration cycles with 3 M guanidine hydrochloride (GuHCl) successfully restored the baseline after each iteration. A representative titration series in buffer for the 6-AH-cAMP derivative is shown in Fig 4A, demonstrating binding responses across all concentrations. Similarly, the titration series in 50% human serum produced sensorgrams comparable in shape and absolute signal to those obtained in buffer, confirming the sensor’s ability to detect PKA-R–cAMP interactions in complex media (Fig 4B). Although minor signal fluctuations were observed in the serum measurements, they were attributed to the instrument’s variable recognition and tracking of the focal spots of the prototype image processing software rather than non-specific binding from serum components. The sensor maintained reliable performance in both conditions, with no evidence of non-specific binding in serum. During these experiments, optimal experimental settings were refined iteratively to achieve the ideal flow rates and best transition times between association and dissociation phase. Throughout these iterations, sample depletion was frequently observed during multiple injections, likely caused by non-specific sticking of PKA-R to the vials and microfluidic system [28]. To mitigate this, *buffer A* was supplemented with bovine serum albumin (BSA) at a concentration of 1 mg/mL, which successfully eliminated sample depletion. While BSA effectively blocked non-specific sticking within the fluidics, its adsorption to the chip surface introduced additional considerations.

**Fig 4.**
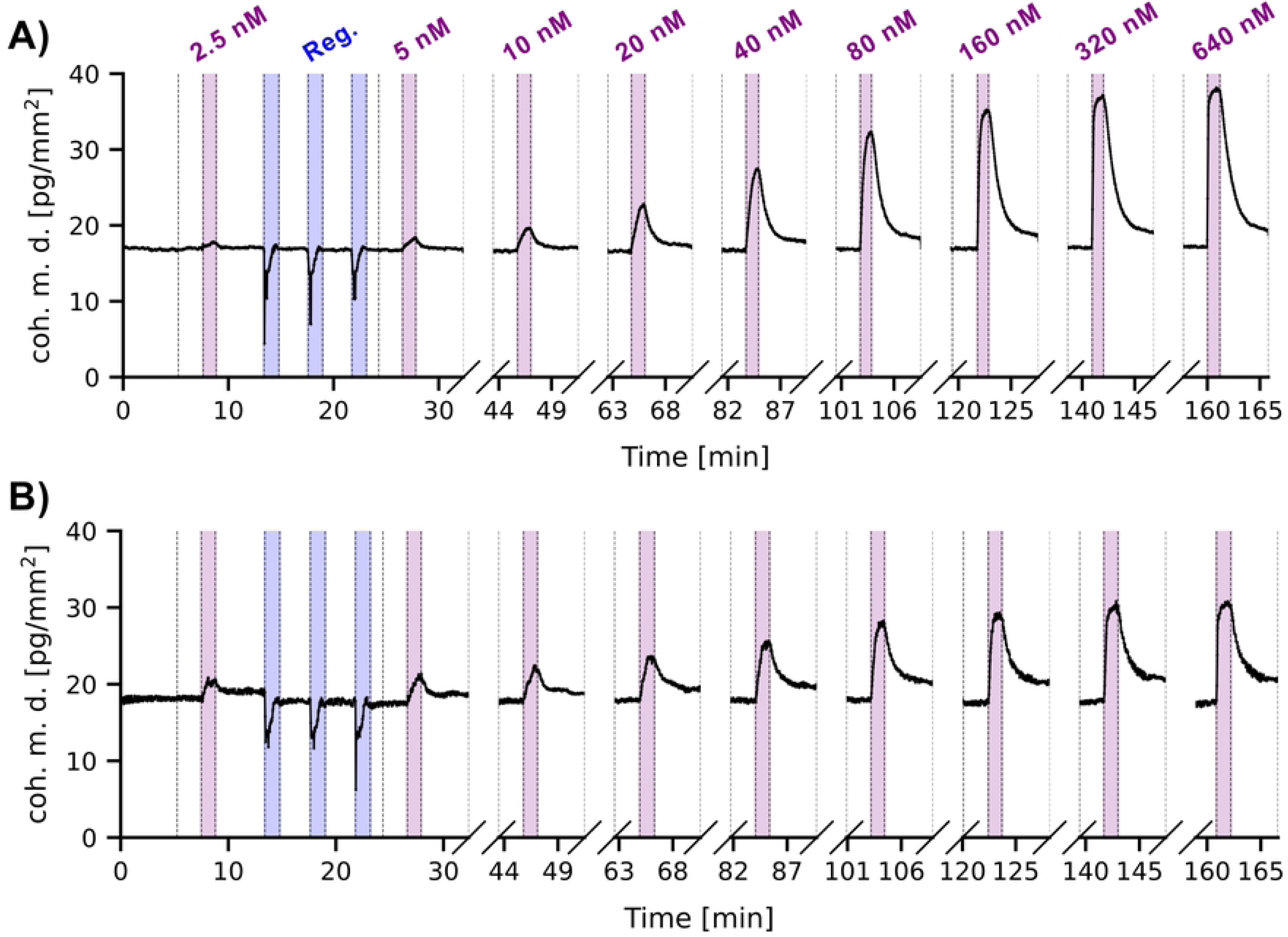
Kinetic experiments in buffer and 50% human serum. **(A)** PKA-R titration conducted in buffer at 20*^◦^*C with 6-AH-cAMP immobilized on the sensor. Binding responses across PKA-R concentrations (2.5 nM to 640 nM, highlighted in pink) demonstrate reproducible sensorgrams. Regeneration cycles with 3 M GuHCl successfully restored the baseline after each cycle. For clarity, only the regeneration step after the first interaction is shown (highlighted in blue); subsequent regeneration cycles have been excluded for better visualization of the full experiment. **(B)** PKA-R titration conducted in 50% human serum at 20*^◦^*C with 6-AH-cAMP immobilized on the sensor with the same experimental procedure as in the buffer experiment in A.

Specifically, the use of 3 M GuHCl for surface regeneration between injections likely caused partial unfolding of the adsorbed BSA, leaving sticky residues. These residuals may have interacted non-specifically with PKA-R, potentially contributing to the incomplete dissociation observed during the dissociation phases. This represents a trade-off between preventing sample loss and maintaining a good sensor chip surface for accurate kinetic measurements [29, 30].

It is important to note that the sensorgrams in buffer and serum do not start at the same coherent mass density or saturate at the same level. This discrepancy arises because each mologram exhibits a slightly different quality, termed analyte efficiency, due to the reactive immersion lithography process [25]. Yet, this does not influence the kinetic parameters as the relative shape of the sensorgrams after proper normalization is independent of the quality of the molograms.

### Kinetic data from focal molography

Fig 5 presents a comprehensive panel of kinetic fits for both cAMP derivatives measured under two different conditions: *buffer A* and 50% human serum. In this panel, each experimental condition is represented by three replicate measurements (labeled as i), ii), and iii)), resulting in a total of 12 kinetic sensorgrams. These sensorgrams, along with the corresponding Langmuir 1:1 binding model fits, demonstrate the reproducibility of the binding kinetics across all replicates. It is important to note that during each experiment, 9 sensorgrams were recorded, according to the flow configuration chosen for these experiments. For the analysis in this paper we only evaluated one sensorgram per experiment as the prototype software of the device produced image registration artefacts (inaccuracies in tracking the focal spots in the image) in most of the sensorgrams, unfortunately.

**Fig 5.**
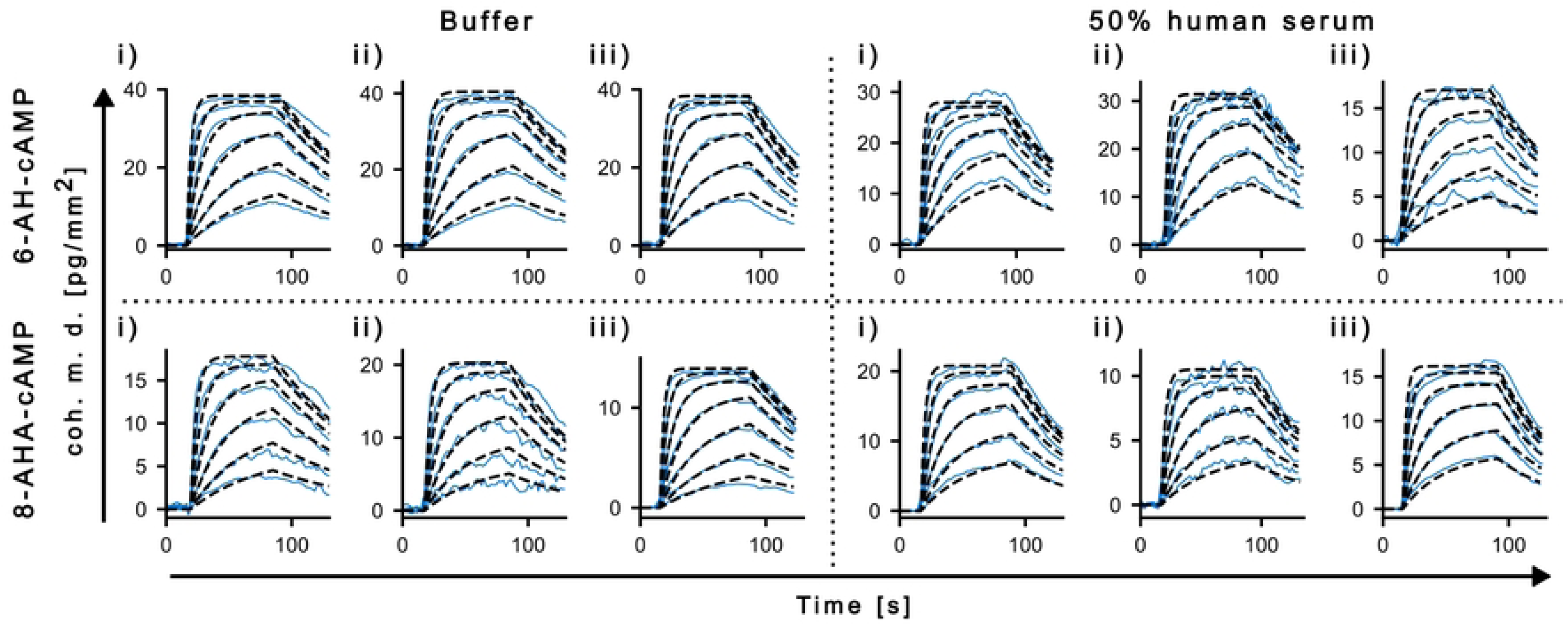
Kinetic fits in buffer and 50% human serum. The panel shows the kinetic fits for both 6-AH-cAMP and 8-AHA-cAMP at 20*^◦^*C under two conditions: *buffer A* and 50% human serum. For each compound and medium, three replicates (i, ii, iii) are displayed. The results demonstrate reproducible binding kinetics (blue lines) and good agreement with the Langmuir 1:1 binding model (dashed black lines).

Overall, the binding responses are consistent in shape in buffer and serum.

Nevertheless, subtle differences in the association rate constant (*k*_on_) and dissociation rate constant (*k*_off_) are evident between the two media. These variations likely stem from the complex biological environment in serum, where interactions with additional serum components may slightly alter the binding dynamics. To ensure that the extracted kinetic parameters accurately represent the specific PKA-R–cAMP interaction, only the initial 15% of the dissociation phase was used to determine *k*_off_, thereby minimizing the influence of incomplete dissociation due to the residual sticking of the protein to the surface. Table 2 summarizes the triplicate kinetic measurements for both 6-AH-cAMP and 8-AHA-cAMP in buffer and serum. The equilibrium dissociation constants (*K*_D_) remain in the nanomolar range under all conditions.

**Table 2.**
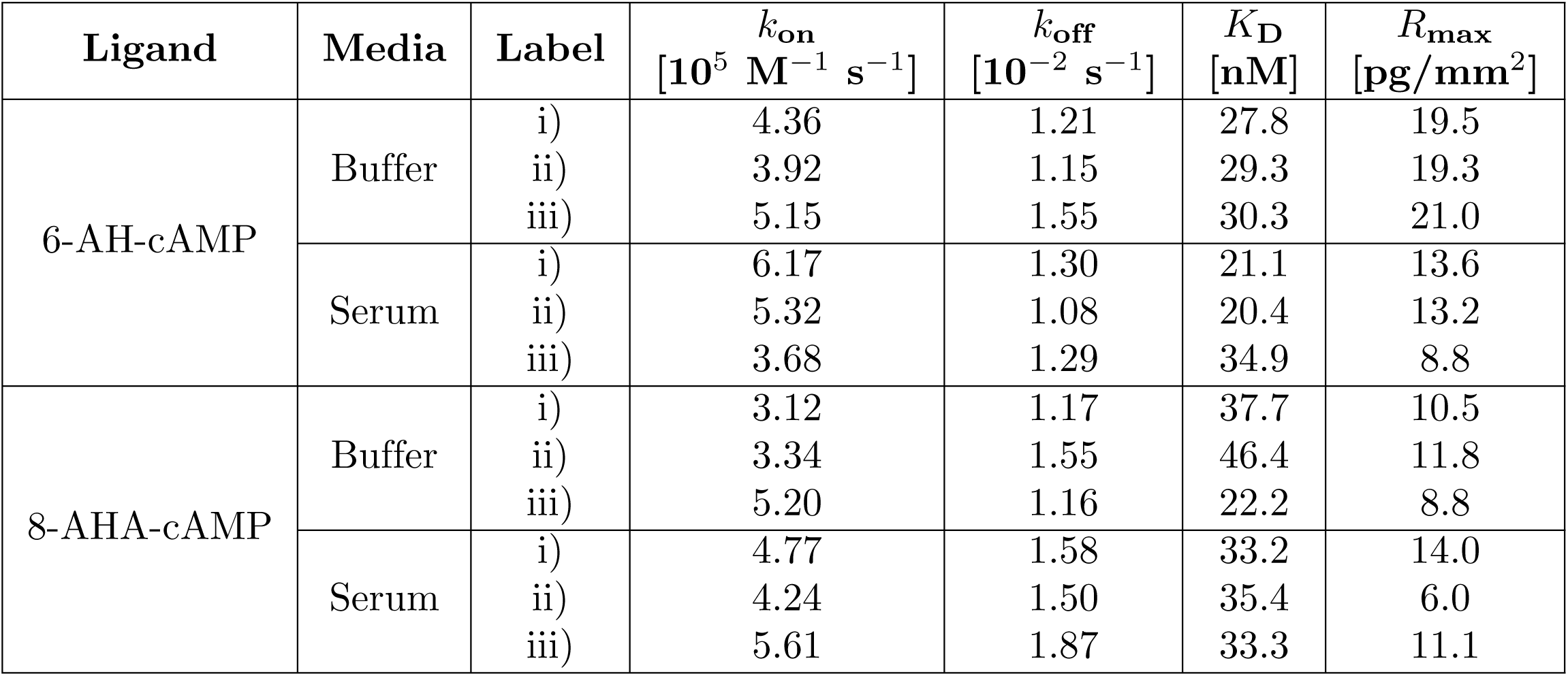
Triplicate kinetic measurements in *buffer A* and 50% serum at 20*^◦^* C for both cAMP derivatives (6-AH-cAMP and 8-AHA-cAMP).

**Table 3.**
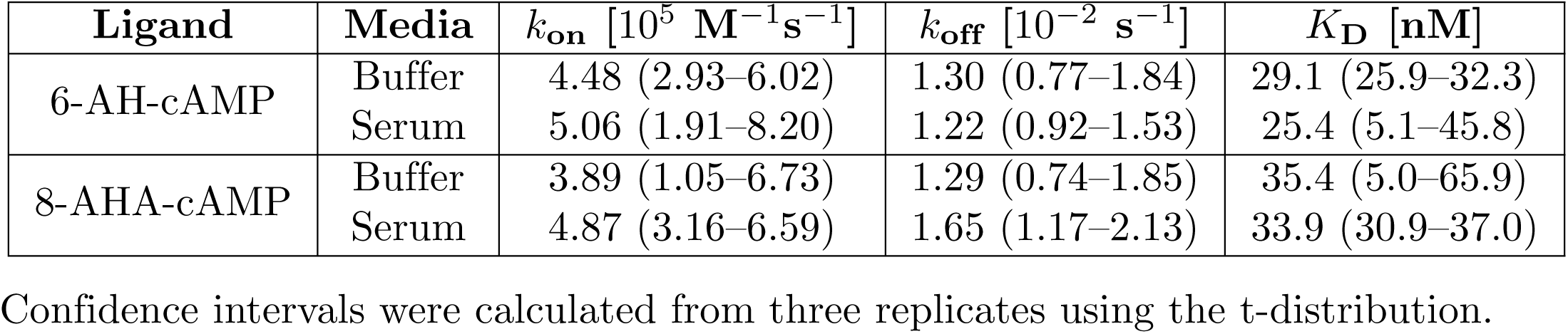
Mean kinetic parameters (*k*_on_, *k*_off_, *K*_D_) with 95% confidence intervals (in parentheses) for 6-AH-cAMP and 8-AHA-cAMP in buffer and 50% serum at 20*^◦^*C.

#### Part 2: Characterization of thermodynamic parameters of interactions with focal molography

Focal molography is not only insensitive to non-specific binding but also remarkably robust against temperature fluctuations. In contrast, temperature ramping on refractometric biosensors, such as SPR, typically requires lengthy equilibration periods—often 30 minutes to an hour—making thermodynamic measurements a low-throughput process. By eliminating the need for such equilibration, focal molography enables faster and more accurate determination of thermodynamic parameters, offering a clear advantage for molecular interaction studies.

#### Temperature fluctuation baseline

To assess the potential impact of temperature fluctuations on focal molography measurements, a temperature ramp experiment was conducted. The sensor was sequentially exposed to controlled 5*^◦^*C increments from 15*^◦^*C to 35*^◦^*C, followed by corresponding decreases back to 15*^◦^*C. During the *Temp. Ramp Up* phase, running buffer (*buffer A*) was flowed over the chip, maintaining a stable baseline even during the ramps. During the temperature plateau phases, dummy injections of the same buffer were performed using the automated sample delivery system of the instrument. Coherent mass density was recorded throughout the experiment, as shown in Fig 6. Although fluctuations and drift in coherent mass density are visible on the graph, it is crucial to note that these deviations are extremely small, occurring within a range of less than 1 pg/mm^2^ (less than 1 RU on an SPR system). These minor variations are well within the noise threshold of the system and can therefore be considered negligible.

**Fig 6.**
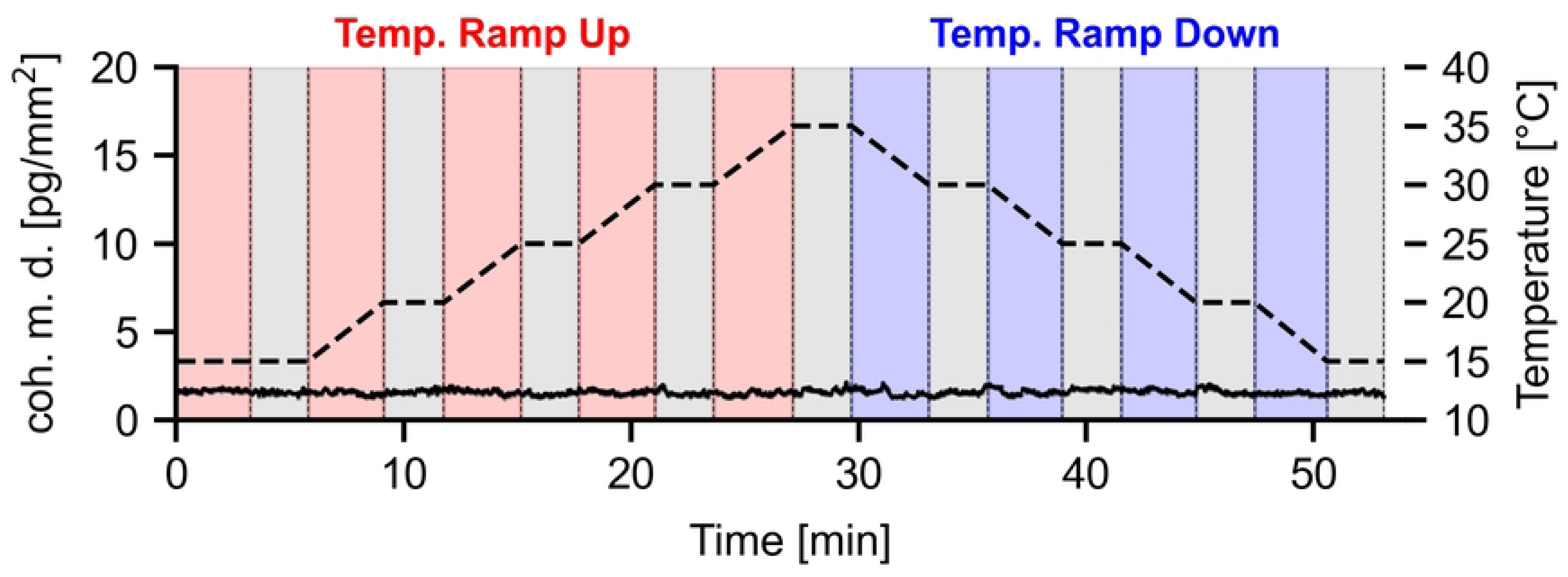
Temperature fluctuation baseline measurement with focal molography. Baseline measurement of coherent mass density (pg/mm^2^) during temperature ramping from 15*^◦^*C to 35*^◦^*C and back to 15*^◦^*C in increments of 5*^◦^*C. Areas shaded in red indicate a temperature increase of 5*^◦^*C and areas shaded in blue indicate a temperature decrease of 5*^◦^*C. Gray shaded areas indicate stable temperature time intervals. The dashed black line represents the applied temperature profile (axis right). Fluctuations in the coherent mass density remain within less than 1 pg/mm^2^, demonstrating system stability under varying temperature conditions.

### Thermodynamic analysis of the interaction of PKA-R with cAMP derivatives via van’t Hoff plots

To investigate the thermodynamic forces governing the PKA-R–cAMP interaction, we measured binding affinities across five temperatures (15–35*^◦^*C) and fit them with a Langmuir 1:1 model (S3 Fig). The resulting equilibrium dissociation constants (*K*_D_) were then used to construct van’t Hoff plots (Fig 7). These plots represent the linear relationship between the natural logarithm of the equilibrium dissociation constant (ln(*K*_D_)) and the inverse temperature (1*/T*), as described by the van’t Hoff equation (eq. 3). From the slope and intercept of the linear fits, we extracted the standard enthalpy (Δ*H*) and entropy (Δ*S*) changes of binding.

**Fig 7.**
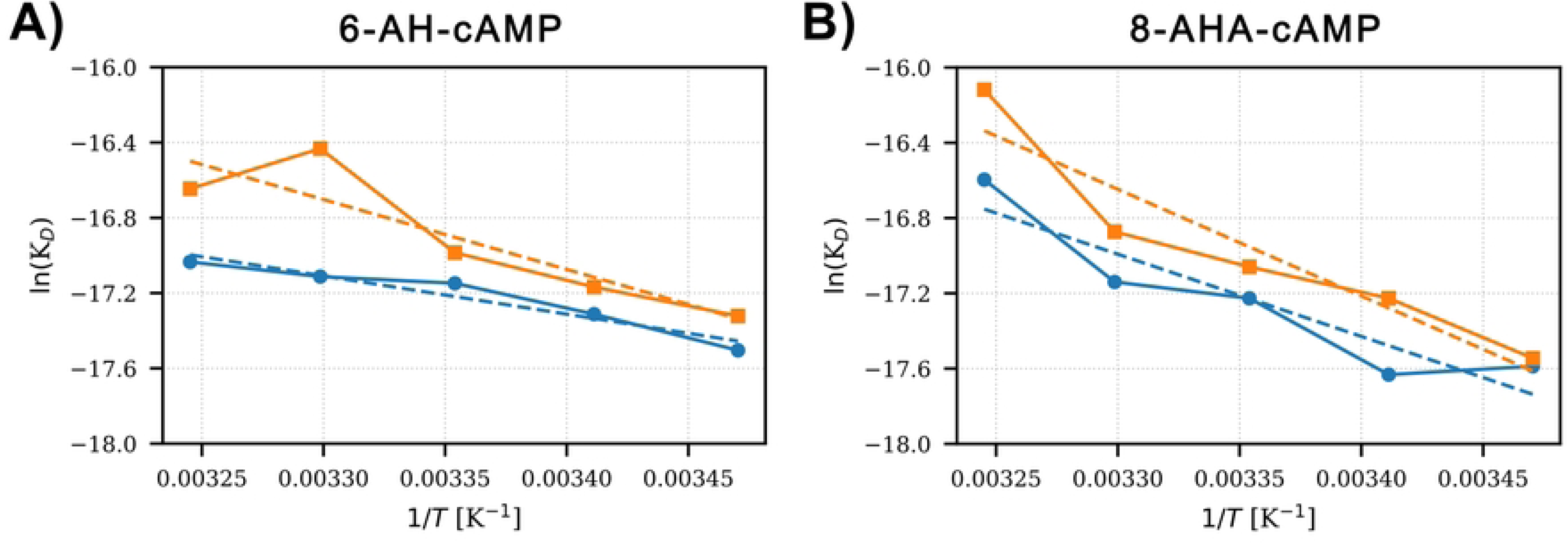
van’t Hoff analysis of temperature-dependent binding affinity. **(A)** 6-AH-cAMP and **(B)** 8-AHA-cAMP. Plots show the natural logarithm of the dissociation constant (ln(*K*_D_)) against the inverse temperature (1*/T*) in buffer (blue) and 50% human serum (orange). Linear fits were used to extract thermodynamic parameters.

Fig 8 summarizes the resulting thermodynamic parameters calculated at 293 K (see S1 Table for data). For both ligands, binding to PKA-R remains highly favorable in both media, with Δ*G* values around −10 kcal/mol yielding similar affinities. However, the underlying driving forces appear to differ between buffer and serum. In buffer, both ligands exhibit favorable entropic contributions, particularly 6-AH-cAMP, which shows predominantly entropy-driven interactions (−*T* Δ*S* = −6.1 kcal/mol) and only moderate enthalpic contribution (Δ*H* = −4.0 kcal/mol). In contrast, the interaction in 50% serum becomes more enthalpy driven for both ligands. For 6-AH-cAMP, Δ*H* drops to −7.4 kcal/mol, while the entropic contribution increases to −2.6 kcal/mol. For 8-AHA-cAMP, the transition is even more striking: binding becomes entirely enthalpy driven in serum, with an unfavorable entropy change (−*T* Δ*S* = 1.2 kcal/mol) and a strong enthalpic contribution (Δ*H* = −11.3 kcal/mol). All thermodynamic parameters were extracted from the linear fits to the van’t Hoff plots via standard error propagation from the covariance matrix returned by the linear regression. Specifically, standard deviations for Δ*H*, Δ*S*, and Δ*G* were calculated using the variances and covariances of the fitted slope and intercept. The reported *K*_D_ values represent the predicted equilibrium dissociation constants at 293 K based on the linear fits, with associated uncertainties derived from the propagated variance in ln(*K*_D_).

**Fig 8.**
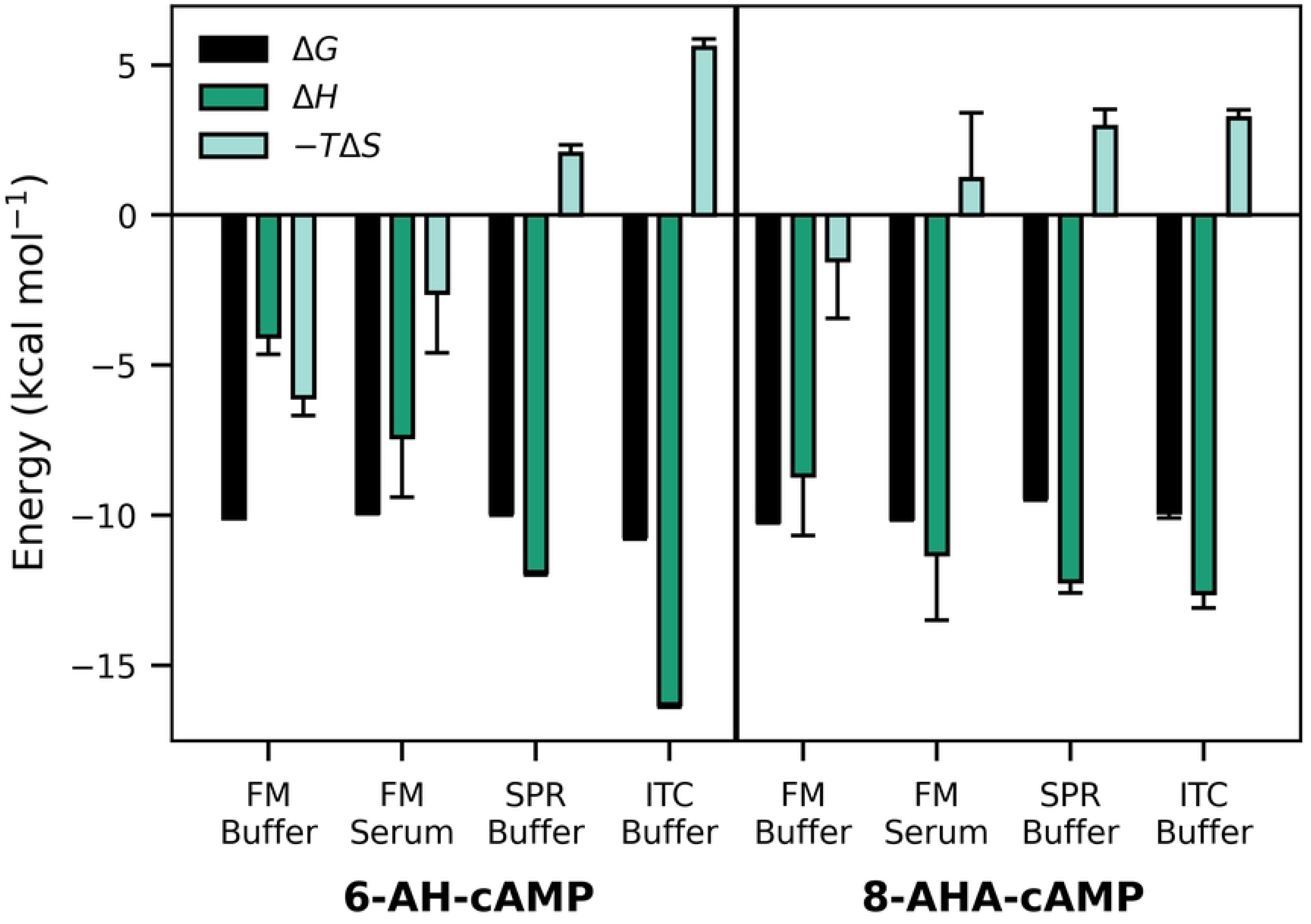
Equilibrium thermodynamic parameters of PKA-R binding at 293. **K.** Grouped bar chart showing Δ*G* (open bar), Δ*H* (hatched bar), and −*T* Δ*S* (solid bar) for 6-AH-cAMP (left block) and 8-AHA-cAMP (right block). Within each ligand, bars are shown for Focal Molography (FM) in buffer and serum, Surface Plasmon Resonance (SPR) in buffer, and Isothermal Calorimetry (ITC) in buffer (Moll et al [20]).

These results suggest that the binding thermodynamics shift considerably in complex media, likely due to altered hydration, specific protein interactions, or changes in molecular flexibility. Notably, entropy contributes less, or even unfavorably, in serum, particularly for 8-AHA-cAMP. While our focal molography measurements allow direct comparison between buffer and serum conditions, the SPR and ITC data from Moll et al. [20] were acquired in buffer only. Comparing across techniques, all three methods yield broadly consistent affinities, yet differ in the magnitude and direction of enthalpic and entropic contributions. These deviations likely stem from methodological differences: ITC captures interactions entirely in solution, while SPR and focal molography rely on surface-immobilized ligands with reduced rotational degree of freedom. Importantly, the two dimensional polyethylene glycol (PEG) based antifouling architecture used in focal molography provides a highly inert and hydrated surface, in contrast to the carboxymethyl-dextran matrix typical of SPR chips [12, 31]. These differences in surface chemistry, ligand presentation, and solvent structuring can influence the thermodynamic fingerprints observed. Thus, while absolute thermodynamic parameters may vary across platforms, such comparisons remain valuable for contextualizing the impact of measurement conditions on biomolecular interactions.

### Transition-state thermodynamics via Eyring analysis

To better understand the energetic barriers behind binding and dissociation, we performed Eyring analyses across the full temperature range. While van’t Hoff analysis uses the temperature dependence of *K*_D_ to determine Δ*H* and Δ*S*, Eyring analysis uses the temperature dependence of association and dissociation rate constants to make statements about the thermodynamics of the transition states of the interaction. Fig 9 shows the resulting plots for both ligands, with buffer and 50% serum data separated into individual sensorgrams for association and dissociation.

**Fig 9.**
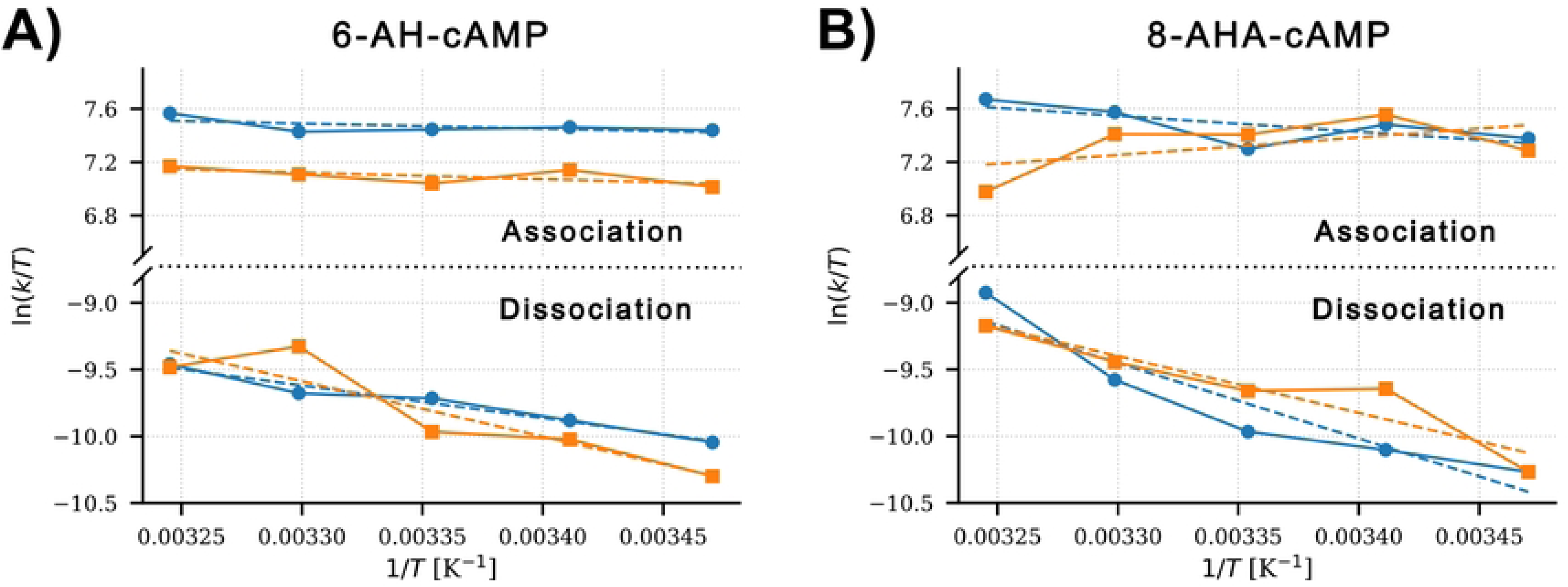
Eyring analysis of PKA-R–cAMP interactions in buffer and 50% human serum. **(A)** 6-AH-cAMP and **(B)** 8-AHA-cAMP. Plots show ln(*k*_on_*/T*) and ln(*k*_off_*/T*) versus inverse temperature (1*/T*) across 15–35*^◦^*C in buffer (blue) and 50% serum (orange). Linear fits enable the calculation of the activation enthalpy (Δ*H^‡^*) and entropy (Δ*S^‡^*) for association and dissociation using the Eyring equation (eq. 5) and from this the derivation of the activation free enthalpy (Δ*G^‡^*).

The plots exhibit approximately linear behavior across the examined temperature range (15–35*^◦^*C) within the experimental error, supporting the applicability of transition state theory. From the slope and intercept of each linear fit, we derived the activation enthalpy (Δ*H^‡^*) and entropy (Δ*S^‡^*) for both association and dissociation steps. These parameters provide insight into the energy barriers and molecular rearrangements required to form or break the PKA-R–cAMP complex. The resulting activation parameters are summarized in Table 4. For 6-AH-cAMP, the association steps in both buffer and serum show low enthalpic barriers (Δ*H^‡^ <* 1kcal/mol) but strongly negative activation entropy, suggesting a transition state that is more ordered than the separated reactants. In contrast, the dissociation step exhibits higher activation enthalpies and even more negative entropy contributions, reflecting either reduced conformational freedom or additional solvent ordering as the complex traverses the dissociation barrier. For 8-AHA-cAMP, a more pronounced difference is observed between conditions. In buffer, association is modestly endothermic (Δ*H^‡^*= 2.38 kcal/mol) and entropically unfavorable, whereas in serum, the enthalpic barrier becomes negative (Δ*H^‡^*= −2.62 kcal/mol), possibly reflecting an enthalpy-favored rearrangement induced by serum components. The dissociation of 8-AHA-cAMP in both media features high enthalpic barriers and strong negative entropy terms, consistent with a stable, tightly bound complex that required significant energetic input to dissociate.

**Table 4.** Transition-state thermodynamic parameters (. Δ*G^‡^*, Δ*H^‡^*, and *T* Δ*S^‡^*) for association and dissociation of PKA-R with 6-AH-cAMP and 8-AHA-cAMP for 293 K in buffer, serum, and previously published SPR data for comparison (Moll et al. [20]).

| Ligand | Process | Condition | $\Delta G^\ddagger$ [kcal/mol] | $\Delta H^\ddagger$ [kcal/mol] | $-T\Delta S^\ddagger$ [kcal/mol] |
| --- | --- | --- | --- | --- | --- |
| 6-AH-cAMP | Association | Buffer | $9.50 \pm 0.8$ | $0.76 \pm 0.6$ | $8.73 \pm 0.56$ |
| | | Serum | $9.72 \pm 0.9$ | $0.98 \pm 0.6$ | $8.74 \pm 0.62$ |
| | | Buffer (SPR) | $9.1 \pm 0.1$ | $6.6 \pm 0.1$ | $2.34 \pm 0.29$ |
| | Dissociation | Buffer | $19.59 \pm 0.7$ | $4.85 \pm 0.5$ | $14.74 \pm 0.50$ |
| | | Serum | $19.69 \pm 2.9$ | $8.25 \pm 2.1$ | $11.45 \pm 2.05$ |
| | | Buffer (SPR) | $19.0 \pm 0.1$ | $18.2 \pm 0.2$ | $0.88 \pm 0.29$ |
| 8-AHA-cAMP | Association | Buffer | $9.52 \pm 1.9$ | $2.38 \pm 1.4$ | $7.14 \pm 1.32$ |
| | | Serum | $9.53 \pm 3.3$ | $-2.62 \pm 2.4$ | $12.14 \pm 2.32$ |
| | | Buffer (SPR) | $9.4 \pm 0.1$ | $6.3 \pm 0.6$ | $3.22 \pm 0.88$ |
| | Dissociation | Buffer | $19.71 \pm 3.3$ | $11.27 \pm 2.3$ | $8.45 \pm 2.28$ |
| | | Serum | $19.59 \pm 2.5$ | $8.47 \pm 1.8$ | $11.12 \pm 1.76$ |
| | | Buffer (SPR) | $18.8 \pm 0.1$ | $18.0 \pm 1.0$ | $0.88 \pm 0.44$ |

Overall, the Eyring analysis suggests that both cAMP derivatives form relatively ordered transition states, with dissociation having the higher activation energies.

Observed differences between buffer and serum conditions may indicate altered transition-state stabilization in complex media, potentially influenced by macromolecular crowding, modified hydration layers, or specific interactions with serum proteins. While not perfectly linear, the trends captured by focal molography provide valuable insights into the energetic landscape of biomolecular interactions under physiologically relevant conditions.

## Discussion

This study demonstrates that focal molography can accurately resolve both equilibrium and transition-state thermodynamics of protein-ligand binding in buffer and complex media. Using the PKA-R–cAMP model, we show that serum alters enthalpy–entropy balance without affecting overall affinity. These results validate focal molography as a robust tool for kinetic and thermodynamic analyses across diverse sample types.

### Overview of results

Focal molography resolved the kinetic and thermodynamic landscape of the PKA-R interaction with 6-AH-cAMP and 8-AHA-cAMP in both buffer and 50% human serum:

- **Kinetics.** Switching from buffer to 50 % serum alters the rate and affinity constants only modestly: *k*_on_ rises by ∼ 13 % for 6-AH-cAMP and ∼ 25 % for 8-AHA-cAMP, *k*_off_ shifts by ∼ −6 % and ∼ +28 %, and the resulting *K*_D_ changes by ∼ −12 % and ∼ −4 %, respectively. All shifts fall within overlapping 95 % confidence intervals, so the differences are not statistically significant.
- **Equilibrium thermodynamics.** van’t-Hoff analysis (Fig 8 and S1 Table) shows two distinct regimes. In buffer, 6-AH-cAMP is rather *entropy-driven* (Δ*H* ≈ −4 kcal mol*^−^*^1^, −*T* Δ*S* ≈ −6 kcal/mol), whereas 8-AHA-cAMP is more *enthalpy-driven* (Δ*H* ≈ −9 kcal/mol, −*T* Δ*S* ≈ −1.5 kcal/mol). Measuring in 50 % serum redirects roughly 3 kcal/mol of free energy from the entropic to the enthalpic term for both ligands, yet Δ*G* shifts by less than 0.2 kcal/mol. The near-constant Δ*G* despite compensating changes in Δ*H* and −*T* Δ*S* exemplifies classical enthalpy–entropy compensation. Compared with SPR and ITC, the molography surface yields a less exothermic Δ*H* (4–12 kcal mol*^−^*^1^ higher) and a negative −*T* Δ*S* (except for 8-AHA-cAMP in serum). This may reflect differences in hydration and crowding between a PEG-based 2D surface and dextran hydrogel or bulk solution.
- **Activation parameters.** For both ligands, getting to the transition state of association is mainly an entropy cost (−*T* Δ*S^‡^*≈ +7 to +12 kcal/mol) and the enthalpy term is small. On dissociation, 6-AH-cAMP is still ruled mostly by entropy, whereas 8-AHA-cAMP relies more on enthalpy in buffer (Δ*H^‡^*∼ 11 kcal/mol) and shifts toward entropy control in serum. Across all conditions, the overall barrier stays roughly the same, about 9.5 kcal/mol for association and 19.6 kcal/mol for dissociation.

Collectively, these results show that the crowded serum environment redistributes enthalpic and entropic contributions without large alterations to either the equilibrium affinity or the kinetic barrier heights, underscoring the robustness of focal molography for quantitative binding analysis in complex media.

### Thermodynamic differences of the PKA-R–cAMP interaction between SPR and focal molography

Although the dissociation constants (*K*_D_) derived from focal molography closely match those obtained by SPR for the PKA-R–cAMP system, the underlying thermodynamic parameters diverge. In buffer, focal molography reveals entropy-dominated binding for 6-AH-cAMP (large −*T* Δ*S* and modestly favorable Δ*H*) and a more enthalpy-driven profile for 8-AHA-cAMP. By contrast, published SPR data report enthalpy-driven binding with positive −*T* Δ*S* values for both ligands [20]. These differences highlight how sensor architecture can influence the apparent thermodynamic decomposition, even when the overall affinity (Δ*G*) remains unchanged.

A likely source of this difference is the physical architecture of the sensor surface. In SPR-based devices, ligands are typically immobilized within a carboxymethyl-dextran hydrogel, which creates a hydrated 3D matrix [11]. Although this environment provides a rather biocompatible, three-dimensional hydrophilic environment, it can introduce heterogeneity, promote rebinding effects, and influence apparent entropy and enthalpy contributions to binding. In contrast, focal molography employs a PEG-based 2D coating, which confines binding to a flat, solvent-exposed interface with lower hydration and reduced steric constraints. Such architectural differences may make focal molography-derived thermodynamic parameters closer to the intrinsic energetics of the interaction in solution.

Nevertheless, 2D formats are not without trade-offs: lower ligand densities can limit sensitivity for small molecule binding studies, and immobilization orientation may still bias results. Thus, neither format should be viewed as categorically superior, underscoring the need for careful contextualization of surface-derived thermodynamic data and, when absolute enthalpy and entropy values are critical, cross-validation with solution-based techniques such as ITC.

### Detailed interpretation of the thermodynamic findings

**Buffer regime** van’t Hoff analysis shows that binding of 6-AH-cAMP in buffer is strongly entropy-driven, with a free energy contribution from entropy of −*T* Δ*S* ≈ −6.1 kcal/mol and a modest enthalpic term of Δ*H* ≈ −4.0 kcal/mol (Fig 8 and S1 Table). This profile suggests that the main driving force for complex formation is solvent reorganization, such as water release upon binding, rather than direct enthalpic contacts.

8-AHA-cAMP follows a similar trend but with a smaller entropic gain (−*T* Δ*S* ≈ −1.5 kcal/mol) and a more favorable enthalpic component (Δ*H* ≈ −8.7 kcal/mol), indicating that fewer water molecules are displaced or that the binding process imposes a stronger conformational restriction on one or both interaction partners. Despite these different decompositions, both ligands converge on similar overall free energies (Δ*G* ≈ −10.1 kcal/mol), a classic example of enthalpy-entropy compensation [32].

Eyring analysis of the association process further supports this picture. For both ligands, reaching the transition state is dominated by a large, unfavorable entropy term (−*T* Δ*S^‡^* ≈ 7 to 9 kcal/mol) negligible enthalpic cost, consistent with a binding mechanism requiring significant ordering—such as desolvation and orientation alignment—before complex formation. Dissociation, on the other hand, shows a clear enthalpic barrier, particularly for 8-AHA-cAMP (Δ*H^‡^*≈ 11 kcal/mol), indicating that specific stabilizing contacts must be broken during complex rupture. These differences highlight that distinct enthalpic and entropic routes can lead to comparable equilibrium affinities for the two ligands.

**Serum regime.** van’t Hoff analysis shows a marked redistribution of free energy contributions compared to buffer. For both ligands, the binding enthalpy becomes more favorable by 3-4 kcal/mol, while the entropy contribution decreases significantly (by roughly 3 kcal/mol), shifting the balance toward enthalpy driven binding. These shifts likely reflect altered hydration and crowding effects in serum, where additional molecular interactions or solvent structuring reduce entropic gains while stabilizing the bound complex enthalpically. Despite these compositional changes, the overall free energy of binding remains nearly unchanged, again highlighting strong enthalpy-entropy compensation in the system. Eyring analysis indicates that the association barrier remains largely entropically controlled in serum. For 6-AH-cAMP, −*T* Δ*S^‡^*≈ 8.7 kcal/mol and Δ*H^‡^*remains close to zero, mirroring buffer conditions. For 8-AHA-cAMP, the entropy penalty becomes slightly larger in serum (12.1 kcal/mol), and the enthalpy term turns weakly favorable (−2.62 kcal/mol), suggesting partial pre-orientation or stabilizing contacts with serum components may lower the energetic cost of binding. Dissociation remains enthalpy controlled for both ligands, with similar activation free energies (Δ*G^‡^* ≈ 19.6 kcal/mol) as in buffer, indicating that the height of the kinetic barrier is preserved despite environmental changes.

Taken together, these findings indicate that the overall kinetic barriers for association and dissociation remain largely unchanged between buffer and serum for both ligands. While 8-AHA-cAMP shows a possible shift in transition state energetics, the uncertainties prevent firm mechanistic conclusions. This underscores the need for caution when interpreting individual enthalpic or entropic activation components.

Importantly, the presented Δ*G^‡^* values across environments, in line with published SPR data, strengthen confidence in the robustness of the focal molography measurements and suggest that environmental effects primarily redistribute enthalpic and entropic contributions without altering the net barrier height.

### Implications for biophysical characterization

Focal molography bridges the long-standing gap between high-quality kinetic data and physiologically relevant sample conditions. This is crucial given that kinetic behavior observed directly in humans/serum can differ markedly from buffer-only measurements [5]. Unlike SPR, which must trade sensitivity for drift correction and requires refractive-index matching, focal molography performs molecular scale self-referencing, where the ridges and grooves of each mologram experience identical bulk effects, so changes in temperature, buffer or serum composition cancel out in real time. This built-in referencing delivers drift-free baselines even during rapid temperature ramps, enabling thermodynamic titrations that would otherwise be met with long equilibration times on refractometric sensors.

At the heart of this stability is the two-dimensional, lithographically defined binding pattern. Because the recognition sites sit in a sparse, planar array rather than in a three-dimensional hydrogel, diffusion is able to keep pace with binding and classical mass-transport artefacts are minimized. Recent work has shown that this layout preserves true solution kinetics for low-affinity antibodies and small proteins while using only microlitre sample volumes and delivering sub-2% inter-assay CVs in matrices as demanding as 50% serum [16, 25]. The pattern is embedded in a PEG brush that resists non-specific adsorption, so crude media can be injected without the baseline drifts that typically accompany high-protein samples. Together, these features give molography a profile that is closer to solution calorimetry in its thermodynamic accuracy, yet it retains the temporal and low sample consumption usually associated with surface methods.

All measurements reported here were performed on a Callisto pre-series prototype, and the platform is still under active development. Continued refinements in hardware, software, and workflow are expected to further streamline its operation, enhance robustness and throughput, and make routine use more straightforward. In this context we see focal molography as a complementary addition to established biophysical methods—especially useful when sample is limited, purification is not possible, matrix effects are central to the question, or rapid temperature profiling required—rather than a replacement.

## Conclusion

Focal molography enables precise kinetic and thermodynamic characterization of biomolecular interactions in both buffer and complex media such as human serum. Using the PKA-R–cAMP system, we demonstrated that this technique reliably determines kinetic rate constants and thermodynamic parameters under near-physiological conditions, maintaining high sensitivity and stability even during temperature-dependent measurements. Our findings reveal that the local environment significantly alters the enthalpic and entropic contributions to binding, underscoring the importance of studying interactions in relevant matrices rather than in buffer alone.

Compared to established techniques like SPR and ITC, focal molography offers the combined benefits of low sample consumption, temperature resilience, and compatibility with crude samples while providing consistent quantitative data. These strengths position it as a robust, complementary tool for investigating protein-ligand interactions under conditions that closely mimic physiological environments, advancing both fundamental research and applications in therapeutic development.

## Supporting information

**S1 Fig.**
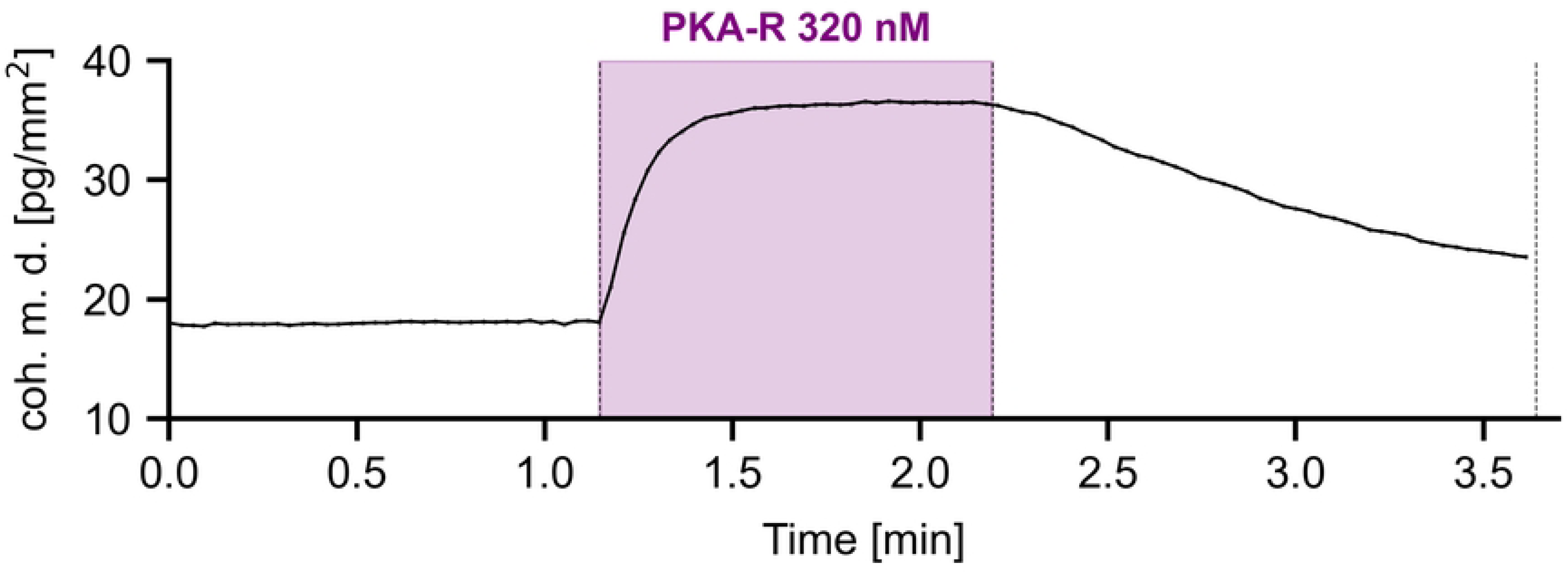
Positive control experiment demonstrating specific binding of PKA-R to immobilized cAMP derivatives. To confirm the effective immobilization of cAMP derivatives on the sensor chip, we conducted a positive control experiment. PKA-R was injected at a concentration of 320 nM in buffer A over the sensor chip surface. This concentration was selected based on a previous study of the same interaction to ensure a detectable signal [20]. As illustrated, the injection of PKA-R resulted in a sharp and substantial increase in coherent mass density (an increase of ≈ 20 pg/mm^2^ within 30 seconds), indicating a successful and specific binding event. This increase stabilized at a higher level, confirming that the free PKA-R molecules effectively interacted with the immobilized cAMP on the sensor surface. These results confirm the sensor’s capability to detect specific PKA-R–cAMP interactions and demonstrate the successful immobilization of cAMP derivatives.

**S2 Fig.**
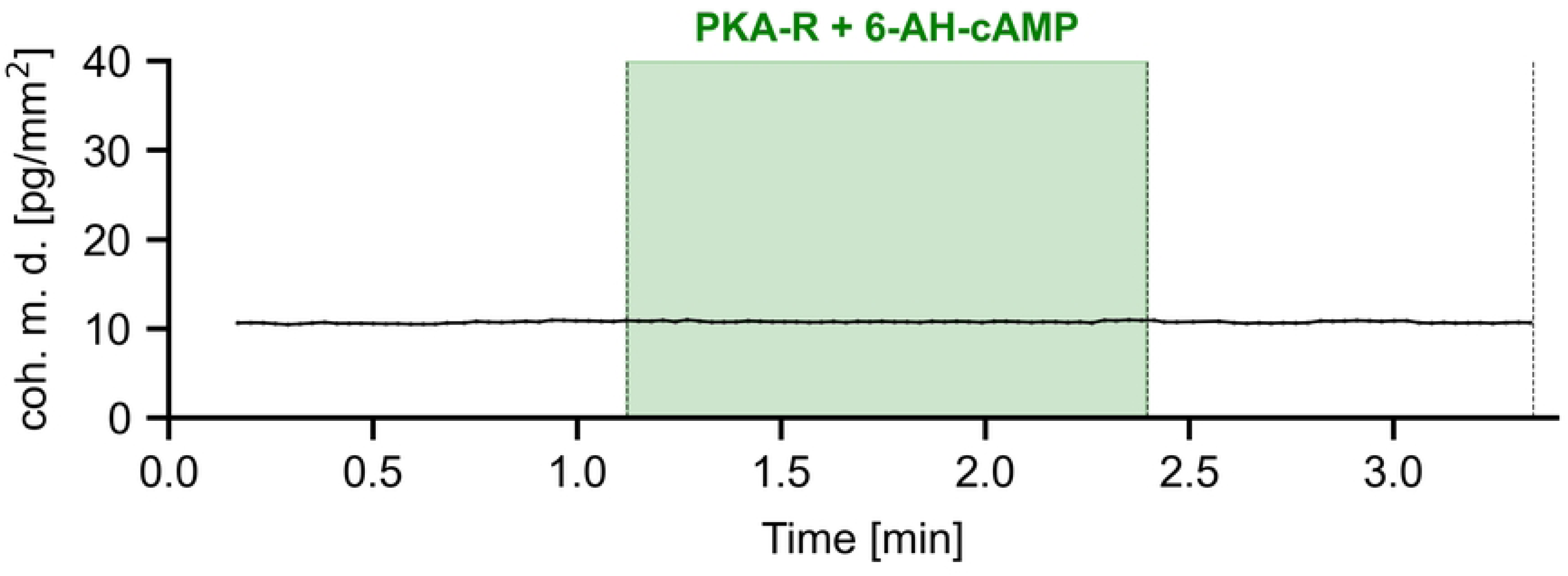
Negative control experiment demonstrating the absence of non-specific binding of PKA-R–cAMP complexes. Following the positive control, a negative control was conducted to verify the specificity of the sensor chip in detecting only unbound PKA-R. In this experiment, a solution containing PKA-R and 6-AH-cAMP was injected in a molar ratio of 1:10 on the sensor surface. This ratio was chosen to ensure that nearly all PKA-R molecules were pre-saturated with 6-AH-cAMP, thereby blocking their binding sites and preventing interaction with the immobilized cAMP on the chip. The injection of this PKA-R–cAMP complex resulted in no significant change in coherent mass density, with only minor fluctuations observed within a range of 0.5 pg/mm^2^. These fluctuations are consistent with baseline noise and well within the acceptable range of stable measurements, indicating that the sensor chip does not detect any changes on the chip surface in the absence of available binding sites on PKA-R. The absence of a binding response in the negative control validates the sensor’s specificity, confirming that only free, unbound PKA-R molecules contribute to the binding signal.

**S3 Fig.**
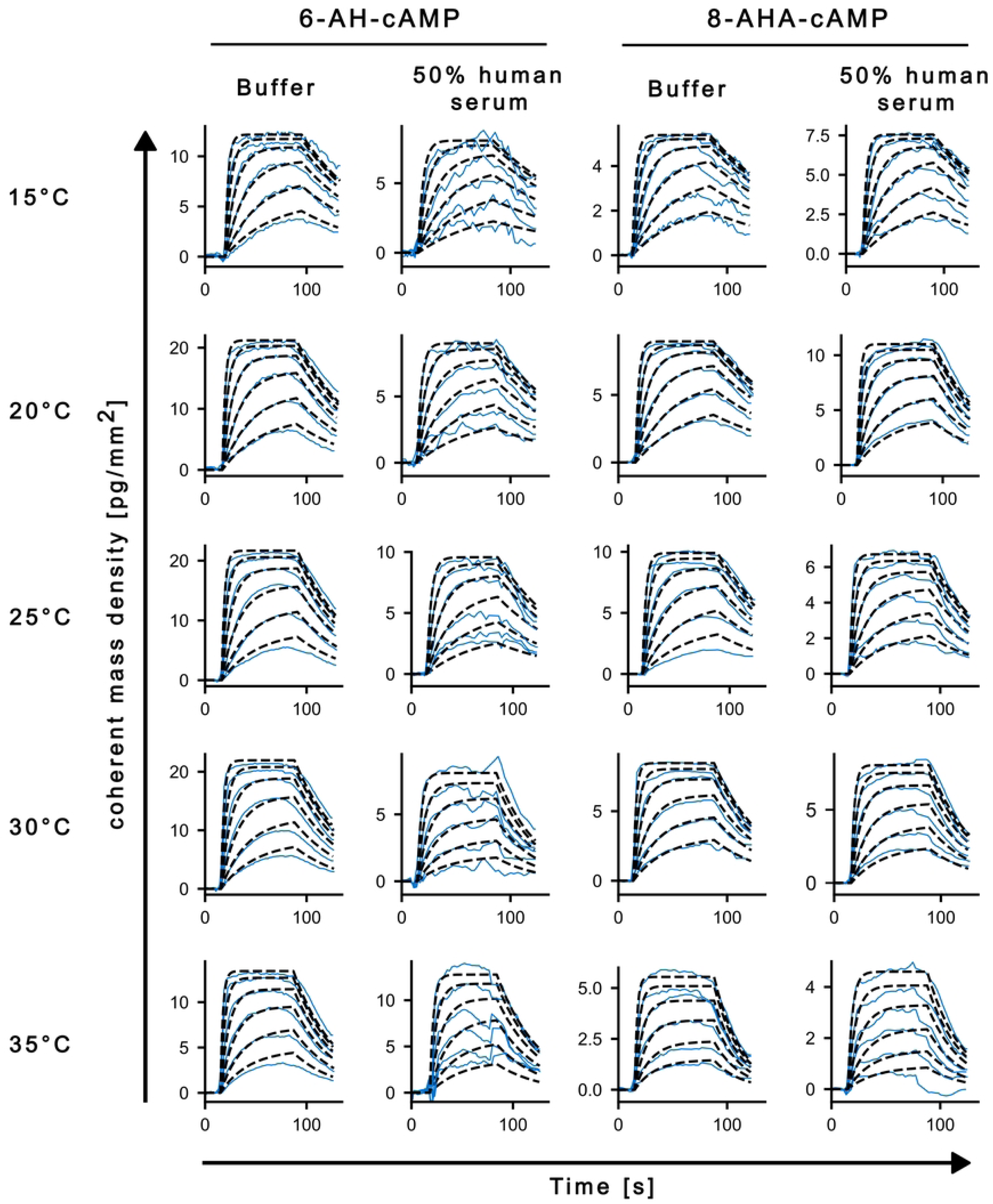
Panel figure of thermodynamic experiment fits. Representative sensorgrams and global kinetic fits for kinetic and thermodynamic analysis. Sensorgrams recorded at 15–35*^◦^*C (5*^◦^*C steps) for the interaction of PKA-R with immobilized 6-AH-cAMP and 8-AHA-cAMP in *buffer A* and 50% human serum. Multi-cycle titrations were globally fit with a Langmuir 1:1 model (dashed black) across all concentrations at each temperature; experimental traces are shown as solid lines. Regeneration cycles are omitted for clarity. Extracted rate constants yielded *K*_D_(*T*) values used for van’t Hoff (Fig 7) and Eyring analysis (Fig 9).

**S1 Table.** Thermodynamic parameters. (*K*_D_, Δ*G*, Δ*H*, *T* Δ*S*) for the interactions of PKA-R with 6-AH-cAMP and 8-AHA-cAMP, calculated for 293 K.

| Ligand | Method | $K_D$ [nM] | $\Delta G$ [kcal/mol] | $\Delta H$ [kcal/mol] | $-T\Delta S$ [kcal/mol] |
| --- | --- | --- | --- | --- | --- |
| 6-AH-cAMP | Focal Mol. (Buffer) | $29.5 \pm 0.9$ | $-10.10 \pm 0.02$ | $-4.04 \pm 0.60$ | $-6.07 \pm 0.62$ |
| | Focal Mol. (Serum) | $36.6 \pm 0.4$ | $-9.90 \pm 0.06$ | $-7.40 \pm 2.00$ | $-2.58 \pm 2.02$ |
| | SPR (Buffer) | $38 \pm 5$ | $-9.9 \pm 0.1$ | $-11.9 \pm 0.1$ | $+2.05 \pm 0.29$ |
| | ITC (Buffer) | $10 \pm 1$ | $-10.7 \pm 0.1$ | $-16.3 \pm 0.1$ | $+5.57 \pm 0.29$ |
| 8-AHA-cAMP | Focal Mol. (Buffer) | $25.5 \pm 2.5$ | $-10.20 \pm 0.06$ | $-8.68 \pm 2.00$ | $-1.50 \pm 1.94$ |
| | Focal Mol. (Serum) | $31.0 \pm 3.4$ | $-10.10 \pm 0.06$ | $-11.30 \pm 2.20$ | $+1.20 \pm 2.20$ |
| | SPR (Buffer) | $110 \pm 21$ | $-9.4 \pm 0.1$ | $-12.2 \pm 0.4$ | $+2.93 \pm 0.59$ |
| | ITC (Buffer) | $46 \pm 15$ | $-9.9 \pm 0.2$ | $-12.6 \pm 0.5$ | $+3.22 \pm 0.29$ |

Focal molography results represent values obtained in this study under both buffer and 50% human serum conditions. SPR and ITC values are reported from Moll et al. (2007) [20], who investigated the same ligand–analyte system. All data are included to facilitate direct comparison between different biophysical techniques. Standard deviations are provided where available.

## Acknowledgments

We gratefully acknowledge the lino Biotech team for their support with laboratory work and data processing, as well as M. Hansch for excellent technical assistance.

